# Advancing Protein Design via Multi-Agent Reinforcement Learning with Pareto-Based Collaborative Optimization

**DOI:** 10.64898/2026.01.13.699365

**Authors:** Mingming Zhu, Jiahua Rao, Xiaoyu Chen, Qianmu Yuan, Yuedong Yang

## Abstract

Protein design is revolutionizing biotechnology, yet existing approaches struggle to balance structural foldability with functional performance. Structure-based models excel at generating stable protein backbones but often overlook critical functional properties, while protein language models capture evolutionary and functional signals but frequently predict sequences lacking structural stability. Integrating these complementary approaches remains challenging due to their inherently conflicting objectives. We present MAProt, a multiagent framework that synergistically combines structure-based and protein language model-based methods for protein design. Each agent specializes in a distinct aspect of the design objective: the structure-based agent (e.g., ProteinMPNN) ensures compatibility with the target backbone, while protein language model-based agents (e.g., ESM, SaProt) capture evolutionary plausibility and functional potential. To reconcile conflicts and achieve optimal trade-offs, we introduce a Pareto-based negotiation module that enables effective multi-objective coordination and consensus among agents. Extensive experiments on benchmark datasets demonstrate that MAProt achieves a remarkable improvement over state-of-the-art baselines, and generalizes robustly across a range of tasks, including thermodynamic folding stability design, functional protein design, and high-affinity antibody design. These results highlight the power of collaborative optimization for advancing rational protein engineering.

**Code:** https://github.com/biomed-AI/MAProt

## Introduction

Protein design is revolutionizing biotechnology by enabling the creation of proteins with customized structural and functional properties (Yeh et al. 2023; Wang et al. 2022). One of the most important challenges is the optimization of protein functional characteristics, which seeks to improve desirable properties, such as stability, binding affinity (Vázquez Torres et al. 2024), or catalytic efficiency (Jiang et al. 2024), through targeted modifications of amino acid sequences. However, the landscape of protein functionality constitutes a vast combinatorial space: as sequence length increases, the number of possible variants grows exponentially, while only a tiny fraction possesses meaningful biological functions (Anishchenko et al. 2021). As design objectives become more sophisticated, achieving both efficient and reliable protein design remains a fundamental challenge.

Despite remarkable advances in protein design methodologies, effectively balancing structural foldability with functional performance continues to be a central obstacle in the field (Notin et al. 2024). Structural stability and functional properties often impose conflicting requirements in the high-dimensional sequence space: mutations that enhance functional performance may disrupt native foldability and stability. Structure-based approaches, such as Protein-MPNN (Dauparas et al. 2022), are highly effective at ensuring structural integrity but often overlook critical functional attributes, including enzymatic activity, binding affinity, or thermostability (Xu et al. 2025). Consequently, these methods may fail to identify variants with the desired functional properties. To address this limitation, recent methods such as ProteinDPO (Widatalla, Rafailov, and Hie 2024) leverage high-quality thermostability datasets to guide structure-based models through reinforcement learning, enabling the generation of sequences that not only preserve the input backbone structure but also exhibit improved functional performance.

However, the scarcity of large-scale, high-quality experimental fitness datasets (Notin et al. 2023) constrains the ability of current models to systematically learn and balance both structural and functional design objectives. In contrast, protein language models such as ESM (Lin et al. 2023) and SaProt (Su et al. 2024) excel at predicting functional effects by capturing global sequence patterns and evolutionary constraints from vast protein databases. This capability enables them to infer functional relevance even in the absence of extensive labeled fitness data (Zhu et al. 2024). Nevertheless, these models frequently generate sequences that lack proper structural stability or foldability. This contrast highlights the complementary strengths of structure-based and language model-based approaches (Yuan et al. 2022), underscoring the need for a unified framework that integrates both structural and sequence information to enable balanced optimization of protein structure and function.

To address this, recent studies (Fei et al. 2025; Xiong et al. 2025) have explored directly integrating structural models with sequence-based language models, aiming to leverage their complementary strengths during protein design. However, negotiating the conflicting objectives of different models within a unified framework remains an exceptionally challenging task (Rao et al. 2022). Simultaneously optimizing multiple objectives, such as foldability, evolutionary plausibility, and functional fitness, inevitably introduces complex trade-offs, as the requirements imposed by different models may be mutually incompatible for certain mutations. Achieving optimal synergy demands careful reconciliation of these competing objectives (Rao et al. 2024). Thus, the central challenge lies not only in integrating heterogeneous predictive signals, but also in effectively resolving inter-model conflicts and establishing consensus among them. Accordingly, there is a pressing need for more robust and principled strategies to negotiate and optimize conflicting objectives within unified protein design frameworks.

In this work, we propose MAProt, a multi-agent-based protein design framework that synergistically combines structure-based and protein language model-based methods. Within this framework, each agent is responsible for a distinct aspect of the design objective: the structure-based agent (ProteinMPNN) ensures compatibility with the target backbone, while sequence-based agents (ESM, SaProt) capture global sequence features and mutational effects, thereby collectively addressing both structural and functional requirements. By formulating protein design as a multi-objective optimization problem, we introduce a Pareto-based negotiation and conflict resolution module that coordinates the potentially competing objectives of different agents and seeks consensus solutions. This enables MAProt to generate protein sequences that optimally balance structural stability and functional potential, providing an effective strategy for the synergistic optimization of protein design.

Extensive experiments on benchmark datasets demonstrate that our method consistently outperforms state-of-theart baselines. MAProt achieves up to 14.1% improvement of success rate in stability protein design on the Megascale benchmark, state-of-the-art performance on the GFP protein fitness optimization benchmark, and a 32% success rate in high-affinity antibody design. Moreover, MAProt is highly extensible to diverse model architectures and parameter scales, demonstrating generalizability across protein domains. Ablation studies and interpretability analyses further underscore the critical role of the negotiation and consensus module in enabling effective multi-objective optimization, establishing MAProt as a powerful and generalizable tool for the rational design and engineering of functional proteins. Our contributions are summarized as follows:

- We propose MAProt, a multi-agent protein design framework that synergistically integrates structure-based models and protein language models.
- We introduce a Pareto-based negotiation mechanism for effective coordination and conflict resolution among agents with competing objectives.
- Extensive experiments on benchmark datasets show that MAProt consistently outperforms state-of-the-art baselines across diverse protein design tasks.

## Related Work

### Protein Design and Optimization

Computational protein design aims to generate amino acid sequences that fold into desired structures and functions. Existing methods are generally divided into two categories. Structure-based models, such as ProteinMPNN and ESM-IF (Hsu et al. 2022), utilize 3D structural information to guide sequence generation, achieving high accuracy in replicating natural protein folds. In contrast, sequence-based methods, including variational autoencoders (Hawkins-Hooker et al. 2021), generative adversarial networks (Repecka et al. 2021), and large language models like ProGen (Madani et al. 2020), learn from evolutionary sequence data to sample diverse and plausible sequences.

Despite these advances, most approaches focus on optimizing general structural or evolutionary fitness, which may not meet specific, application-driven functional requirements. Preference alignment strategies have emerged to guide design toward desired properties; for example, ProteinDPO uses direct preference optimization, while triplet-based distillation (Karimi et al. 2025) and DRAKES (Wang et al. 2024) employ reward-driven fine-tuning. However, these methods depend on high-quality datasets for particular properties and still lack effective mechanisms to integrate and balance these conflicting objectives in protein design.

### Multi-objective Collaborative Optimization

Multi-objective optimization in protein design seeks to simultaneously satisfy diverse criteria, such as foldability, evolutionary plausibility, and functional fitness. Recent methods attempt to leverage both complementary strengths of structure- and sequence-based models (Misaki et al. 2025). For example, del Alamo et al. (2025) improves ProteinMPNN for antibody CDR design by integrating the antibody-specific language model AbLang (Olsen, Moal, and Deane 2022), enhancing native-likeness and binding. AICE (Fei et al. 2025) combines structural and evolutionary constraints with inverse folding to predict high-fitness mutations, while ProteinGuide (Xiong et al. 2025) enables guided generation by conditioning ProteinMPNN and ESM3 (Hayes et al. 2025) on auxiliary signals.

Existing frameworks typically lack principled strategies for conflict resolution, often relying on static or heuristic integration. However, directly optimizing multiple objectives simultaneously inevitably introduces complex trade-offs, as competing requirements may be mutually incompatible. To address this challenge, we introduce a Pareto-based multiagent framework that enables systematic negotiation and consensus-building among heterogeneous models, thereby achieving balanced and synergistic optimization across multiple protein design objectives.

## Methodology

### Problem Formulation

We formulate protein sequence design as an optimization problem, which seeks to find a sequence *S ∈ V*^*L*^, where *V* is the set of standard amino acids and *L* is the sequence length, that maximizes a specific fitness metric *F* = *f* (*S, B*). Here, *B* denotes a protein backbone structure, and the fitness function *f* quantifies desired protein properties.

Our framework leverages a set of specialized pretrained models as agents: ProteinMPNN for structure-based sequence generation, ESM for functional evaluation, and SaProt for assessing sequence and structural constraints. Each agent focuses on a distinct aspect of protein design, either generating sequences or evaluating them based on structural, functional, or evolutionary criteria. Through Pareto-based optimization, these agents negotiate to balance multiple objectives and reach consensus on high-quality sequences. This multi-agent strategy enables MAProt to efficiently explore the protein fitness landscape and generate sequences with superior biochemical properties.

### Architectures of ProteinMPNN, ESM and SaProt

To leverage the diverse information in protein structures and sequences, MAProt employs three specialized agents, each tailored to a distinct modality of protein data. Specifically, ProteinMPNN leverages structural information by utilizing the 3D backbone coordinates, which are represented as a tensor in ℝ^*L*×*3*^, where *L* denotes the sequence length. This allows ProteinMPNN to generate amino acid sequences *S* = (*s*_1_, …, *s*_*L*_) that are compatible with the target fold.

ESM is a transformer-based protein language model that processes amino acid sequences *S* = (*s*_1_, …, *s*_*L*_), where *s*_*i*_ denotes the residue at position *i* (Yuan et al. 2024). By capturing long-range dependencies, ESM evaluates the evolutionary plausibility and functional potential of candidate sequences based on patterns learned from large-scale protein sequence datasets. In contrast, SaProt integrates both sequence and structural information by encoding the 3D structure as a structural sequence *F* = (*f*_1_, …, *f*_*L*_) using Foldseek (Van Kempen et al. 2024), where each *f*_*i*_ denotes the structural token at residue *i*. These structure tokens are then concatenated with the amino acid sequence *Z* = ((*s*_1_, *f*_1_), …, (*s*_*L*_, *f*_*L*_)), enabling joint modeling of sequence-structure relationships. Further details are provided in Appendix.

### Single-Agent Preference Alignment

We next analyze how each agent, operating independently on its respective modality, develops distinct sequence design preferences, establishing a foundation for subsequent multiagent integration and consensus.

#### Building Preference Data

To align sequence generation preferences with experimentally measured protein fitness, we construct a set of sequence preference pairs, following prior work ProteinDPO: 𝒟_pref_ = (*x, y*^*w*^, *y*^*l*^), where for each input *x, y*^*w*^ and *y*^*l*^ are candidate sequences with higher and lower experimentally measured fitness, respectively. These pairs are independently generated for each agent using available experimental data, ensuring that preference comparisons are empirically grounded.

#### Direct Preference Optimization

Each agent is initialized from a pretrained model, trained on large-scale sequence or structure data, to model the conditional distribution *y ~ π*_*θ*_(*·*|*x*), where *θ* denotes model parameters. To align sequence generation with experimental fitness, we fine-tune each agent’s policy to maximize the likelihood of high-fitness sequences. Specifically, we apply Direct Preference Optimization (DPO) (Rafailov et al. 2023) to align the policy *π*_*θ*_ with the preference dataset 𝒟_pref_, relative to a fixed reference policy *π*_ref_.

DPO defines the probability that a candidate sequence *y*_*w*_ is preferred over another candidate *y*_*l*_ in prompt *x* as:

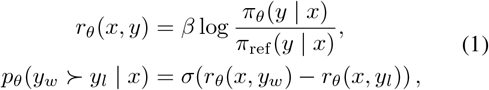

where *σ*(*·*) denotes the sigmoid function and *β* is a temperature hyperparameter. The DPO loss for each agent is:

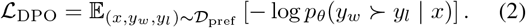

To fine-tune generation preferences toward proteins with desired properties, we jointly minimize the sum of the DPO losses for ProteinMPNN, ESM, and SaProt:

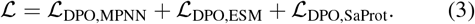

The preference alignment mechanism fine-tunes each model with DPO to align its generative preferences with its specialized property, such as structural compatibility, evolutionary relevance, or functional potential.

### Pareto-based Multi-Agent Preference Agreement

While each agent can independently optimize its sequence generation policy, the protein design often requires integrating multiple, potentially conflicting objectives. To this end, we propose a multi-agent consensus module, inspired by the concept of Pareto optimality in multi-objective optimization, to systematically quantify and leverage inter-agent agreement for robust preference optimization. In this framework, consensus regions correspond to Pareto-optimal solutions that balance multiple desirable properties, thereby reflecting agreement across diverse agent preferences.

For each native protein *x* in the training set, we generate *N* sequences 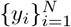 using the policy model learned from the previous stage. Each agent independently evaluates every sequence, obtaining three probability scores:

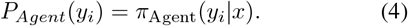

Due to the distinct modeling strategies and domain expertise of each agent, these probabilities can exhibit significant variability. Therefore, a systematic approach is needed to identify candidates with high inter-agent consensus.

#### Consensus-Guided Preference Pair Construction

To quantify consensus and conflict between two agents, we compute the relative preference margin: 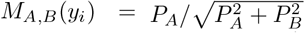 where *P*_*A*_ and *P*_*B*_ are the probability scores assigned by agents *A* and *B* to candidate *y*_*i*_. We assign a consensus label *g*_(*A,B*)_(*y*_*i*_) to each candidate sequence as follows:

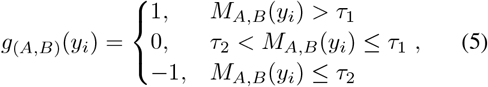

where *g*_(*A,B*)_ = 0 indicates a consensus region where both agents provide similar assessments, while *g*_(*A,B*)_ = 1 or *−*1 indicates a conflict region, reflecting a significant preference for one candidate by either agent.

Candidates *y*_*i*_ with the same group label are grouped, reflecting consistent evaluation regimes across agent pairs. Preference pairs *D*_*cons*_ = (*x, y*^*w*^, *y*^*l*^) are constructed within each group, ensuring *y*^*w*^ and *y*^*l*^ keep preference consistency.

#### Consensus Detection

To robustly evaluate the reliability of each constructed preference pair (*y*^*w*^, *y*^*l*^), we combine structural and continuous consensus measures. Specifically, let *p* = *y*^*w*^ *− y*^*l*^ denote the vector of probability score differences across agents, with the ideal consensus direction given by *u* = (1, 1, 1). The continuous consensus is measured by the angular deviation *α* between *p* and *u*:

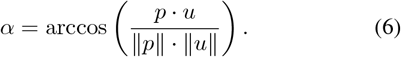

For the structural consensus, we calculate the proportion of consensus labels *g* = 0 within each group, denoted as *a*, which reflects the level of agreement among agents. These two components are integrated to form the consensus score:

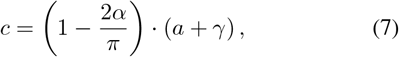

where *γ* controls the baseline confidence level, *π* is circle constant pi.

#### Consensus-Weighted Loss Formulation

Finally, we apply this consensus score as a weighting factor, encouraging the model to prioritize learning from high-confidence pairs and to be less influenced by uncertain or disputed ones. The confidence-weighted loss is:

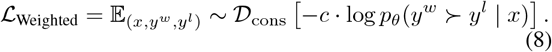

### Multi-Agent Sampling

To leverage the strengths of multiple agents, we employ a logit fusion strategy during sequence generation. At each residue position *i*, we aggregate information from all *M* agents by summing their logits to obtain the fused logits:

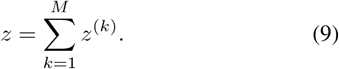

The probability distribution over amino acid types is:

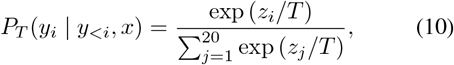

where *T* is a temperature hyperparameter. At each position, the next residue is sampled from the distribution with multinomial sampling, enabling stochastic sequence generation.

## Experiments

### Experimental Setup

#### Datasets

We evaluate MAProt on three benchmarks: Megascale (Tsuboyama et al. 2023) (*~* 1.8M variants) for stability optimization, GFP (Sarkisyan et al. 2016) (56,806 variants) for functional phenotype design, and AffinityDesign (47 complexes) for high-affinity antibody–antigen engineering. Each dataset targets a distinct protein property. Further details and data splits are provided in the Appendix.

#### Baselines

We compare our method against a comprehensive set of baselines covering diverse protein sequence optimization strategies. On the Megascale and AffinityDesign datasets, we include guidance-based methods such as classifier guidance (CG) (Nisonoff et al. 2025), SMC-based guidance and TDS (Wu et al. 2023); classifier-free guidance (CFG) (Ho and Salimans 2022); pretrained models without task-specific fine-tuning, ProteinMPNN and ESM-IF; as well as reinforcement learning fine-tuned methods, DRAKES, and ProteinDPO. On the GFP dataset, we benchmark against a range of optimization and sampling baselines, including GFlowNets (GFN-AL) (Jain et al. 2022), model-based adaptive sampling (CbAS) (Brookes, Park, and Listgarten 2019), greedy search (AdaLead) (Sinai et al. 2020), Bayesian optimization (BO-qEI) (Wilson et al. 2017), conservative model-based optimization (CoMS) (Trabucco et al. 2021), proximal exploration (PEX) (Ren et al. 2022), and predictor-guided approaches such as VLGPO (Bogensperger et al. 2025), GWG, and GGS (Kirjner et al. 2024).

#### Experimental Settings

MAProt is implemented using Pytorch v1.10.2 and trained on a single NVIDIA A800 80GB GPU. All trainable parameters are optimized by Adam (Kingma and Ba 2014) algorithm with a learning rate of 5 *×* 10^*−*6^. We determined the DPO loss hyperparameter *β* and temperature *T* with grid search. For all baselines, they are retrained on the same machine with the same hyperparameter settings reported. Details on hyperparameters and evaluation metrics are in the supplementary materials.

### Main Results

#### Thermodynamic Folding Stability Design

We evaluate our method on the challenging task of protein thermodynamic folding stability design using the Megascale benchmark. Following previous work, we evaluate all methods using the ΔΔ*G* predictor provided by DRAKES. Table 1 summarizes the performance of MAProt and baseline methods. Our method achieves state-of-the-art performance improving success rate by +14.1% and median predicted ΔΔ*G* by +43.1% over DRAKES, demonstrating superior generation of stable protein sequences.

**Table 1:** Comparison of the performance of baselines and MAProt on the Megascale benchmark for protein stability optimization. Results are reported as the mean across 3 random seeds, with standard deviations in parentheses.

| Method | Pred-ddG(Med)↑ | %(ddG > 0)(%)↑ | scRMSD(Med)↓ | %(scRMSD < 2)(%)↑ | Success Rate(%)↑ |
| --- | --- | --- | --- | --- | --- |
| ESM-IF | -1.898(0.011) | 2.9(0.1) | 1.105(0.021) | 73.5(0.5) | 2.0(0.1) |
| ProteinMPNN | -0.510(0.001) | 35.1(0.0) | 0.862(0.027) | 93.0(0.4) | 31.5(0.1) |
| CG | -0.561(0.045) | 36.9(1.1) | 0.839(0.012) | 90.9(0.6) | 34.7(0.9) |
| SMC | 0.659(0.044) | 68.5(3.1) | 0.841(0.006) | 93.8(0.4) | 63.6(4.0) |
| TDS | 0.674(0.086) | 68.2(2.4) | **0.834(0.001)** | 94.4(1.2) | 62.9(2.8) |
| CFG | -1.186(0.035) | 11.0(0.4) | 3.146(0.062) | 29.4(1.0) | 1.3(0.4) |
| DRAKES | 1.095(0.026) | 86.4(0.2) | 0.918(0.006) | 91.8(0.5) | 78.6(0.7) |
| ProteinDPO | 1.120(0.078) | 89.8(4.9) | 1.036(0.034) | 82.9(0.6) | 73.7(4.7) |
| **MAProt (DPO)** | **2.143(0.030)** | **93.8(0.5)** | 1.012(0.038) | 89.0(2.3) | 82.9(2.7) |
| **MAProt** | **1.603(0.073)** | **94.7(1.2)** | 0.890(0.032) | **94.8(1.7)** | **89.7(0.2)** |

MAProt robustly optimizes folding stability while maintaining structural integrity. DRAKES preserves structure but has limited exploration, yielding smaller ΔΔ*G* gains. Multi-agent frameworks without consensus (e.g., MAProt (DPO)) can improve ΔΔ*G*, but often at the cost of structural quality due to conflicting objectives. In contrast, MAProt’s consensus-driven negotiation effectively balances evolutionary and structural constraints, delivering superior stability without sacrificing integrity, demonstrating the advantage of our multi-agent consensus optimization approach.

To further demonstrate the effectiveness of MAProt, we analyze the distribution of predicted ΔΔ*G* values for generated sequences (Figure 2). MAProt’s distribution is shifted toward higher stability, with a substantially larger proportion of successful designs than other methods. For example, the designed sequence for 4G3O closely aligns with the original backbone (scRMSD = 0.245) while achieving significantly improved stability, also indicating that MAProt enhances stability without compromising structural integrity.

**Figure 1.**
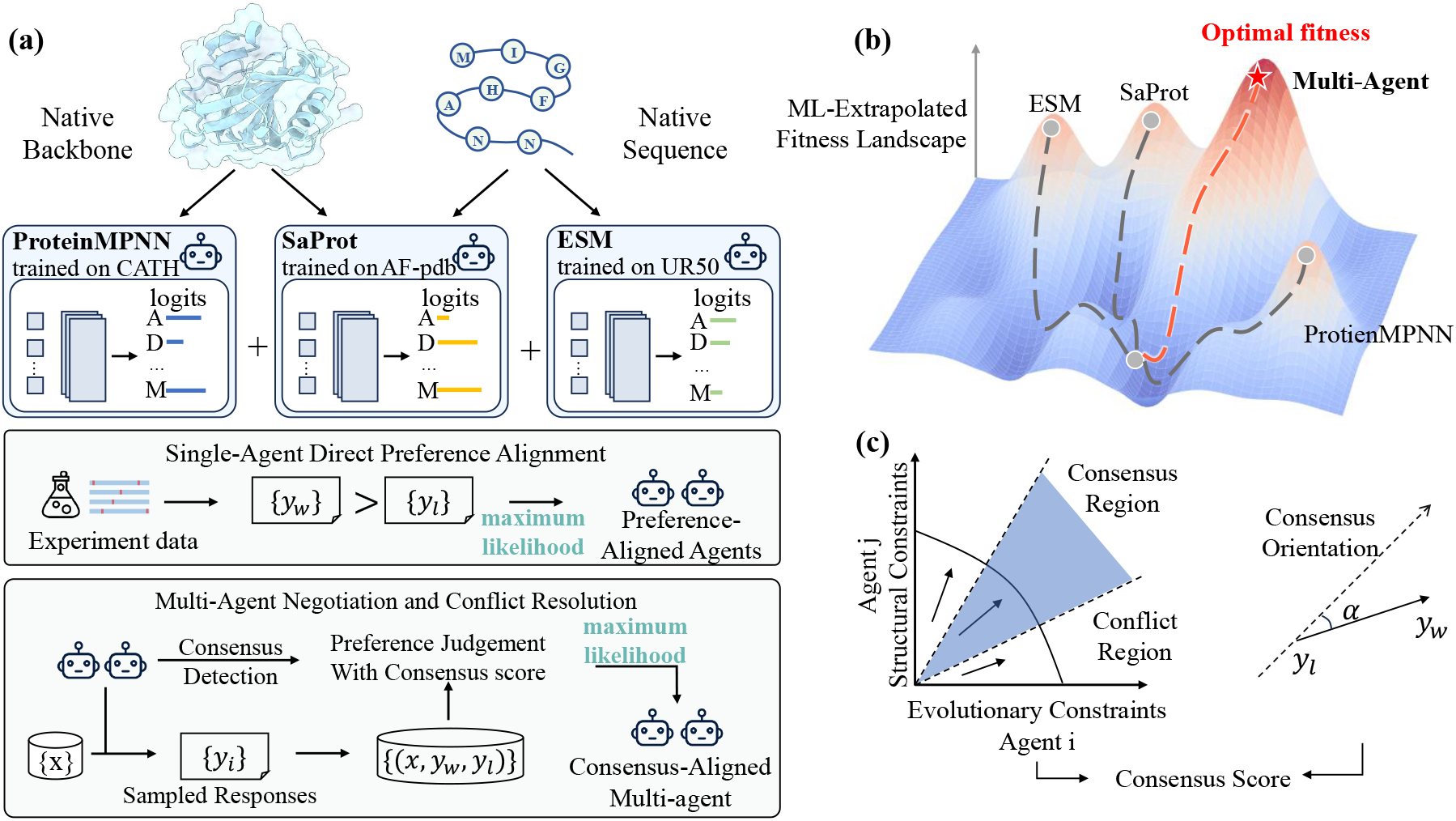
An overview of the MAProt framework. (a) MAProt integrates multiple sequence design models—ProteinMPNN, SaProt, and ESM. Experimental data is first used to align the preferences of individual agents. Subsequently, multi-agent negotiation, incorporating consensus detection and conflict resolution, yields a consensus-aligned solution. (b) Agents collaboratively explore an extrapolated fitness landscape to identify optimal sequences. (c) The consensus and conflict regions among agents are visualized in terms of structural or evolutionary constraints, with a consensus score quantifying agreement.

**Figure 2.**
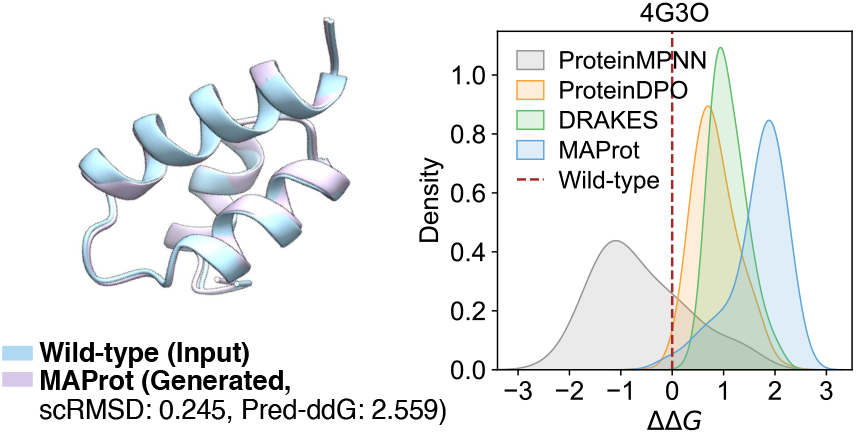
Example of generated proteins. The histograms display the ΔΔ*G* for each generated sequence.

#### Functional Protein Design: Green Fluorescent Protein

We further evaluate MAProt on the challenging task of functional protein design using the GFP benchmark. Unlike stability optimization, this task requires models to generalize to novel, high-fluorescence sequences across a vast and diverse sequence space.

Table 2 presents the results on the GFP benchmark, comparing MAProt with baselines across both medium and hard difficulty tasks. MAProt consistently outperforms all baselines in the primary objective, achieving mean fitness scores of 0.88 (medium) and 0.87 (hard), establishing a state-of-the-art performance. The improvement is particularly pronounced on the hard task, where MAProt achieves up to 12% higher fitness than the second-best method (VLGPO with 0.78), demonstrating robust generalization and optimization. In addition, MAProt maintains competitive diversity and novelty scores, indicating that generated sequences are not only highly functional but also sufficiently distinct. This superior performance can be attributed to MAProt’s ability to effectively coordinate multi-agent preferences, balancing functional optimization with sequence diversity. By contrast, single-agent or less coordinated approaches tend to converge to local optima or produce redundant variants, as reflected in their lower novelty and diversity metrics.

**Table 2:**
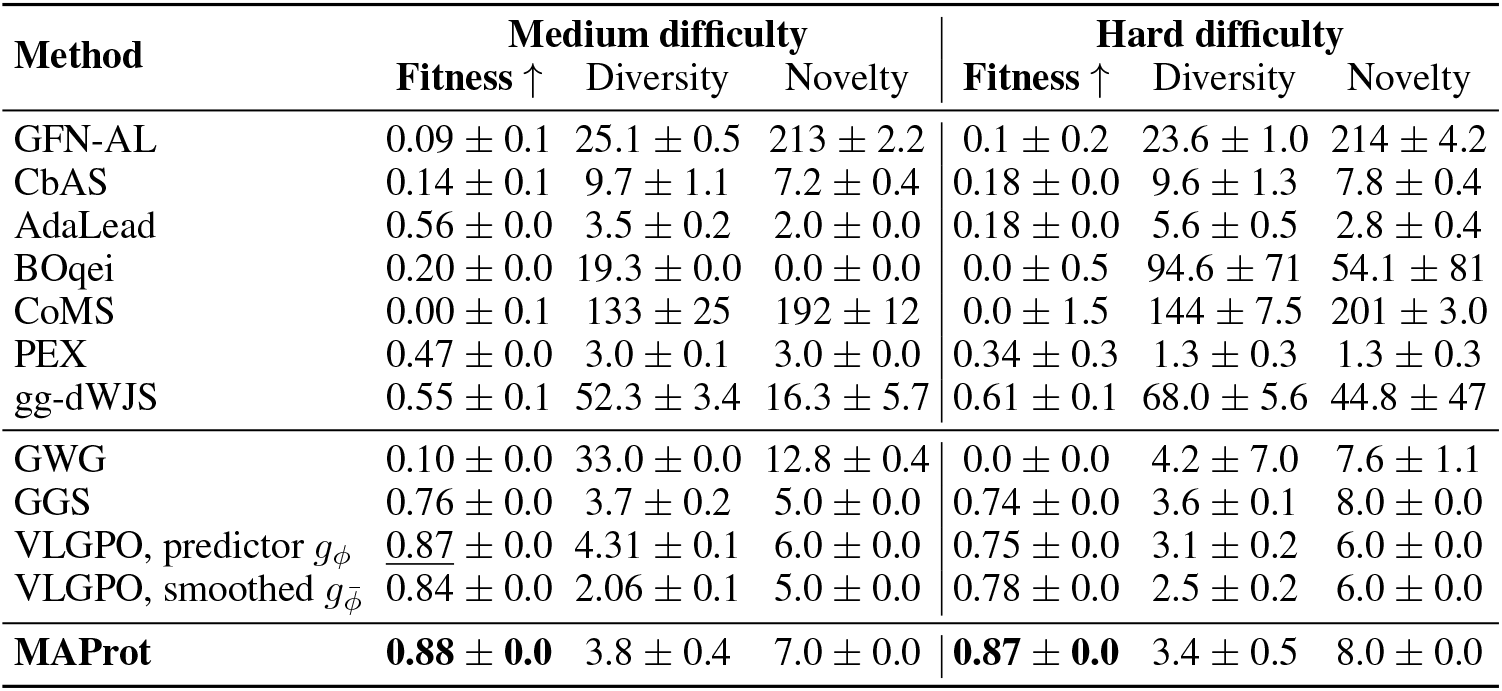
Comparison of the performance of baselines and MAProt on the GFP benchmark for fluorescence fitness optimization. Results are reported as the mean across 5 random seeds, with standard deviations in parentheses.

#### High-Affinity Antibody Design

To further demonstrate the practicality and robustness of MAProt, we apply it to high-affinity antibody sequence design, a task requiring simultaneous optimization of multi-chain complexes for structural compatibility and binding affinity, closely reflecting real-world therapeutic antibody engineering challenges. We benchmark MAProt on the AffinityDesign dataset against state-of-the-art baselines (Table 3). MAProt achieves a 32.1% success rate, a high sequence recovery rate (0.763), and strong binding affinity (mean Δ affinity of 0.153), significantly outperforming all competitors. Compared to the best baseline MAProt (DPO), MAProt achieves a 43.9% higher success rate and further reduces mean Δ affinity (lower is better), demonstrating superior performance on both key metrics. These results highlight MAProt’s strength in integrating multiple agent perspectives—including structure, sequence, and function—when navigating the complex antibody sequence space.

**Table 3:** Comparison of the performance of baselines and MAProt on the AffinityDesign benchmark for antibody affinity optimization. Results are reported as the mean across 3 random seeds, with standard deviations in parentheses.

| Method | ΔAffinity(Med)↓ | $\%(\Delta$Affinity$< 0)(\%) \uparrow$ | Recovery(Med)↑ | $\%($Recovery$> 0.7)(\%) \uparrow$ | Success($\%) \uparrow$ |
| --- | --- | --- | --- | --- | --- |
| ProteinMPNN | 0.889(0.006) | 3.3(0.1) | 0.543(0.000) | 0.0(0.0) | 0.0(0.0) |
| ProteinMPNN(SFT) | 0.342(0.102) | 27.4(2.8) | 0.478(0.005) | 0.0(0.0) | 0.0(0.0) |
| ProteinDPO | 0.345(0.027) | **34.3(0.9)** | 0.542(0.001) | 0.0(0.0) | 0.0(0.0) |
| MAProt (Zero-shot) | 0.301(0.002) | 23.7(0.1) | 0.708(0.002) | 65.0(0.2) | 14.2(0.1) |
| MAProt (DPO) | 0.222(0.010) | 31.8(0.6) | 0.706(0.000) | 58.8(0.5) | 22.3(0.9) |
| **MAProt** | **0.153(0.020)** | **32.4(2.6)** | **0.763(0.001)** | **98.8(0.1)** | **32.1(2.5)** |

### Ablation Study

To assess the contribution of each component in MAProt, we conduct ablation experiments on the Megascale dataset.

#### Multi-Agent Collaboration Optimization

As shown in Table 4, the multi-agent framework consistently outperforms single-agent approaches. With DPO, it achieves a success rate of 82.9% versus 79.1% for the single-agent baseline. For SFT, the multi-agent setting reaches 69.8% compared to 66.7%. This shows that collaborative optimization leads to more robust and diverse sequence generation. With full integration, MAProt achieves the highest success rate of 89.7%, surpassing both the DPO-only multi-agent model (82.9%) and dual-agent settings (e.g., ProteinMPNN+SaProt). These results highlight that combining diverse agent knowledge and advanced strategies in MAProt substantially improves protein design performance over standard approaches.

**Table 4:** Ablation of key components and training strategies on Megascale. +ESM/+SaProt denotes inclusion of the respective language model agents.

| Method | Training | Success Rate(%)↑ |
| --- | --- | --- |
| ProteinMPNN | Zero-shot | 31.5(0.1) |
| MAProt | Zero-shot | 54.4(0.9) |
| ProteinMPNN | SFT | 66.7(3.5) |
| MAProt | SFT | 69.8(1.3) |
| ProteinMPNN | DPO | 79.1(3.5) |
| ProteinMPNN+ESM | DPO | 82.2(3.7) |
| ProteinMPNN+SaProt | DPO | 84.1(2.2) |
| MAProt | DPO | 82.9(2.7) |
| **MAProt** | **Consensus** | **89.7(0.2)** |

#### Hyper-Parameter Analysis

We further analyze the impact of the preference sharpness parameter *β* in multi-agent learning. As summarized in Table S3 (see Appendix for details), low *β* values (e.g., 0.1) enforce strong preference adherence but cause large structural deviations, while high *β* (1.0) better preserves structure but reduces fitness and success rate. An intermediate setting (*β* = 0.5) achieves the best balance, yielding the highest success rate (89.7%) and robust performance. These results suggest that moderate preference strength is crucial for effective alignment without compromising structural quality.

### Visualization Analysis

As shown in Figure 3, Ramachandran plots for MAProt and MAProt (DPO) generated structures are concentrated within favored regions, closely matching native proteins. This confirms that generated structures are geometrically plausible and free from steric clashes. MAProt (top row, orange) shows a more focused distribution in the most favored regions, indicating higher structural stability and fidelity to native conformations. It also covers a broader range of allowed regions than MAProt (DPO), demonstrating greater conformational diversity. Notably, MAProt avoids significant populations in disallowed regions, confirming the absence of geometric conflicts, while MAProt (DPO) shows some residues in these unfavorable areas.

**Figure 3.**
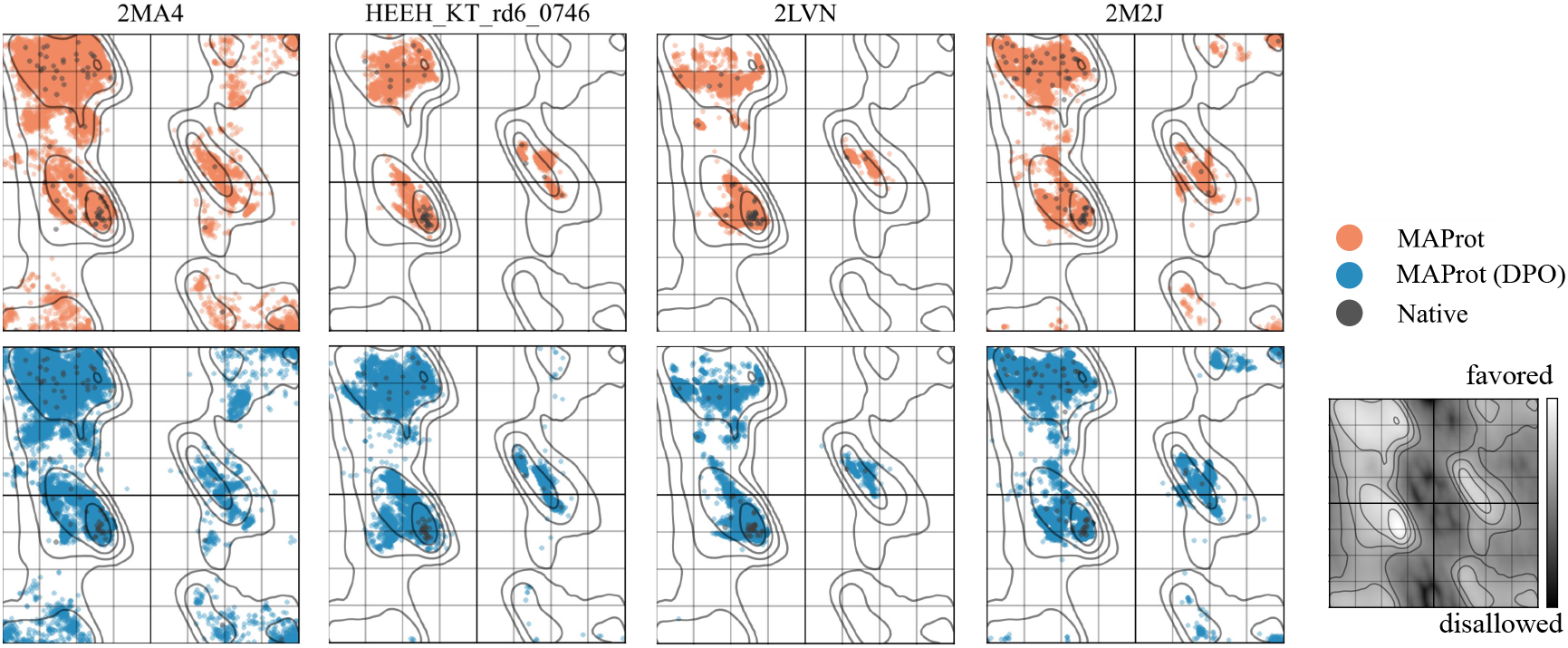
Ramachandran plots comparing dihedral angle *ψ* and *ϕ* distributions for generated versus native proteins; inset shows favored (light) and disallowed (dark) regions.

## Conclusion

In this work, we present MAProt, a flexible and robust multiagent protein design framework that unifies diverse structural and functional objectives through negotiation and consensus. Extensive experiments show that MAProt consistently outperforms state-of-the-art methods across multiple benchmarks, achieving higher success rates and superior functional properties, as further supported by ablation and visualization analyses. In the future, we would like to extend MAProt to enable joint structure and sequence co-design, and apply it to more complex protein assemblies and de novo design tasks. Moreover, integrating experimental feedback and active learning will further enhance MAProt’s practical impact, supporting broader applications in protein engineering and therapeutic discovery.

## Acknowledgments

This study has been supported by the Guangdong S&T Program [2024B1111140001], the China Postdoctoral Science Foundation [2025M771540, GZB20250391], the Youth S&T Talent Support Program of GDSTA [SKXRC2025177], the Young S&T Talent Support Program of Guangdong Precision Medicine Application Association [YSTTGDPMAA2025], and the Lingang Laboratory [LGL-8888].

